# The white matter network in cognitive normal elderly predict the rate of cognitive decline

**DOI:** 10.1101/2020.04.16.044479

**Authors:** Weijie Huang, Ni Shu

## Abstract

White matter degradation has been proposed as one possible explanation for age-related cognitive decline. The human brain is, however, a network and it may be more appropriate to relate cognitive functions to properties of the network rather than specific brain regions. Cognitive domains were measured annually (mean follow-up = 1.25 ± 0.61 years), including processing speed, memory, language, visuospatial, and executive functions. Diffusion tensor imaging was performed at baseline in 90 clinically normal older adults (aged 54–86). We report on graph theory-based analyses of diffusion tensor imaging tract-derived connectivity. The machine learning approach was used to predict the rate of cognitive decline from white matter connectivity data. The reduced efficacy of white matter networks could predict the performance of these cognitive domains except memory. The predicted scores were significantly correlated with the real scores. For the local regions for predicting the cognitive changes, the right precuneus, left inferior parietal lobe and cuneus are the most important regions for predicting monthly change of executive function; some left partial and occipital regions are the most important for the changed of attention; the right frontal and temporal regions are the most important for the changed of language. Our findings suggested that the global white matter connectivity characteristics are the valuable predictive index for the longitudinal cognitive decline. For the first time, topological efficiency of white matter connectivity maps which related to special domains of cognitive decline in the elderly are identified.

## Introduction

In healthy aging, the age-related declines occur in many cognitive functions (Craik and Salthouse, 2008, Craik, FIM.; Salthouse, TA., editors. The handbook of aging and cognition. 3rd ed. Psychology Press; New York: 2008.Li He,CAR 2014). Impairment in multiple domains, especially memory and executive function, predicts more rapid progression to dementia (Gross et al., 2012, cortical signatures of cognition and their relationship to Alzheimer’s disease.; Knopman et al., 2015, Spectrum of cognition short of dementia: Framingham Heart Study and Mayo Clinic Study of Aging.)., Prediction of age-related cognitive decline is needed to differentiate normal aging processes from disease courses that call for intervention.

Disrupted brain white matter communication pathways could be one underlying mechanism for cognitive declines. Diffusion tensor imaging (DTI) is ideally suited for the study of cortical disconnection as it provides indices of structural integrity within interconnected neural networks. It also has the potential to capture broad inter-regional white matter connectivity properties that give rise to successful cognitive function, and thus is a promising candidate predictor of future cognitive function declines(Scott et al., 2017) (Bennett & Madden, 2014a). In the cognitively normal subjects, reduced WM microstructure is linked to poorer cognitive performance, and generally find stronger relationships between white matter microstructure and cognitive function(Madden et al., 2012). Progression of reduced WM tracts predict cognitive decline, with the most pronounced effects on processing speed and executive functions (Jokinen et al., 2011; Kramer et al., 2007; Prins et al., 2005; van Dijk et al., 2008). Meanwhile, other studies found the WM tracts predict the episodic memory over time, but not executive function (Fjell et al., 2016; Rabin et al., 2018). The potential difficulties in interpreting these findings include the selection of white matter tracts examined and the specific tests used to assess the different cognitive domains. However, data are lacking on the association between the brain’s connectivity patterns and cognitive abilities in generally healthy non-demented older participants

The widespread degeneration of WM connectivity between brain regions makes network analysis an appropriate way to explore the possible neuropathological mechanisms of cognitive aging at the system level. The whole WM networks exhibit several topological properties, such as global and local network efficiency, small-worldness and highly connected hub regions(Hagmann et al., 2008; van den Heuvel & Sporns, 2011). With aging, there were changes in the underlying network organization that resulted in decreased global network efficiency(**Gong,Wen,zhao**)(Gong et al., 2009; Perry et al., 2015; Wen et al., 2011; Wiseman et al., 2017; Zhao et al., 2015) (Gong et al., 2009; Perry et al., 2015; Wen et al., 2011; Wiseman et al., 2017; Zhao et al., 2015), hub connections(Gong et al., 2009; Perry et al., 2015; Wen et al., 2011; Wiseman et al., 2017; Zhao et al., 2015) and network strength(Gong et al., 2009; Perry et al., 2015; Wen et al., 2011; Wiseman et al., 2017; Zhao et al., 2015) . Age was found to have a negative effect association with regional efficiency of many cortical regions in elderly person, including most of the frontal and temporal cortical areas and the entire cingulate cortex (Gong et al., 2009; Perry et al., 2015; Wen et al., 2011; Wiseman et al., 2017; Zhao et al., 2015).The topological efficiency was associated with cognitive functions, especially executive functions and processing speed (Lo et al., 2010) (Reijmer et al., 2013) (Shu et al., 2012) (Gong et al., 2009; Perry et al., 2015; Wen et al., 2011; Wiseman et al., 2017; Zhao et al., 2015) (Gong et al., 2009; Perry et al., 2015; Wen et al., 2011; Wiseman et al., 2017; Zhao et al., 2015). Correlations between connectivity of specific regions and cognitive assessments were also observed, e.g., stronger connectivity in regions such as superior frontal gyrus and posterior cingulate cortex were associated with better executive function(Gong et al., 2009; Perry et al., 2015; Wen et al., 2011; Wiseman et al., 2017; Zhao et al., 2015). However, there is limited study about the association between WM network topological properties and the progression of cognitive aging. The predictive value of WM network for cognitive aging in healthy controls remains unknown.

One challenge is that the majority of studies examining the relationship between white matter diffusion characteristics and cognition have been cross-sectional in design. An advantage of longitudinal designs is that within-person change can be directly examined. The few longitudinal studies addressing this issue in clinically normal older adults have had relatively short follow-up periods (Charlton et al. 2010; Lövdén et al. 2014; Ritchie et al. 2015; Köhncke et al. 2016). Studies with longer and more frequent follow-up periods are essential to better assess the influence of diffusion characteristics on cognitive decline in aging. (Rabin et al., CC2018). Another challenge is the statistical analysis method. The prediction exemplifies a larger trend in neuroscience (Bzdok, 2016; Gabrieli et al., 2015; Pereira et al., 2009; Varoquaux & Thirion, 2014) and psychology (Yarkoni & Westfall, 2016) to move from correlative to predictive studies, often using tools from machine learning(Liem F et al. 2017). The machine-learning methods enable predictions on a single-subject level and have been devoted to the prediction of behavior and brain age based on neuroimaging data (Predicting brain-age from multimodal imaging data captures cognitive impairment). The machine learning in cognitive neuroscience has the potential to answer fundamental questions about cognitive functions(Arbabshirani, Plis, Sui, & Calhoun, 2017; La Corte et al., 2016; Liem et al., 2017; Yun et al., 2013). Such an approach has already proven its validity in recent investigations of neuropsychological features in neurology or psychiatry (Costafredaetal., 2011, Pattern of neural responsest over balfluency shows diagnostic specificity for schizophrenia and bipolardisorder.; Quintanaetal., 2012, Using artificial neural networks in clinical neuropsychology: highper for mancein mild cognitive impairment). The model is built using the input feature vectors (e.g., the features of white matter networks) and matching this vector with expected outputs(e.g., prediction of cognitive variables).

Therefore we assessed whether global or regional topological efficiency of WM network predicts cognitive function declines in older adults. Furthermore, we simultaneously plan to clearly identify the predictive value of WM network on each cognitive function domains. We model the effects of WM network topological efficiency on annual change in neuropsychological performance within BABRI data set by machine learning approach. We hypothesized that WM network connectome parameters would add predictive value to cognitive declines in particular in executive functions.

## Method and materials

### Participants

The present sample consisted of 90 clinically normal, community-dwelling older adults from the Beijing Ageing Brain Rejuvenation Initiative (BABRI), which is an ongoing, longitudinal study investigating aging and cognitive impairment in urban elderly people in Beijing, China. At study entry, all participants had a 25 or greater on the Mini-Mental State Examination (MMSE; Folstein et al. 1975, Mini-mental state: a practical method for grading the cognitive state of patients for the clinician. J Psychiatr Res. 12:189–198.), performed within age-adjusted norms on a battery of neuropsychological tests and ?? on the Activities of Daily Living (ADL) scale(Zhang M, Elena Y, He Y. Activities of daily living scale. Neuroimage. 1995;7(suppl 3):289-297). They took part in at least once follow-up neuropsychological test and the interval time of follow up is more than six months. All participants provided a written informed consent for our protocol that was approved by the ethics committee of the State Key Laboratory of Cognitive Neuroscience and Learning at Beijing Normal University. Participants were all right-handed and native Chinese speakers.

### Neuropsychological tests

All the participants received filled in an Activities of Daily Living (ADL) scale and received a battery of neuropsychological tests assessing general mental status and other cognitive domains, such as episodic memory, executive function and language ability. The general mental status of the participants was assessed using the MMSE. The memory tests also included the Auditory Verbal Learning Test (AVLT) (Schmidt, 1996), Digit Span (Gong, 1992), and Rey-Osterrieth Complex Figure test (ROCF) (recall) (Rey, 1941). Participants underwent multiple visuospatial tests, including the ROCF (copy) and the Clock-Drawing Test (Rouleau *et al.*, 1992). The attention function of participants was measured by the Stroop Test (Golden, 1976) and Symbol Digit Modalities Test (SDMT) (Sheridan *et al.*, 2006). Executive function was assessed with the Stroop Test and TMTba test. Lastly, language ability was assessed with the Boston Naming Test (BNT) (Kaplan *et al.*, 1983).

### MRI data acquisition

MRI data were acquired at a 3.0 T Siemens Tim Trio MRI scanner in the Imaging Center for Brain Research at Beijing Normal University. These data included one high-resolution T1 scan and two sets of diffusion magnetic resonance imaging (dMRI) data scans. A high-resolution T1-weighted image covering the whole brain was acquired using sagittal 3D magnetization prepared rapid gradient echo (MP-RAGE) sequences. The scanning parameters included repetition time (TR) = 1900 ms, echo time (TE) = 3.44 ms, inversion time (TI) = 900 ms, slice thickness = 1 mm, no inter-slice gap, 176 axial slices, matrix size = 256 × 256, field of view (FOV) = 256 × 256 mm^2^, flip angle = 9°, and voxel size = 1 × 1 × 1 mm^3^. For each dMRI scan, diffusion images covering the whole brain were acquired by a single-shot echo planar imaging-based sequence using scan parameters consisting of a TR = 9500 ms, TE = 92 ms, 30 diffusion-weighted directions with b-value of 1000 s/mm^2^ and a single image with b-value of 0 s/mm^2^, slice thickness = 2 mm, no inter-slice gap, 70 axial slices, matrix size = 128 × 124, field of view (FOV) = 256 × 248 mm^2^, and voxel size = 2 × 2 × 2 mm^3^.

### Data preprocessing

The preprocessing involved eddy current and head motion correction, estimation of the diffusion tensor and calculation of the fractional anisotropy (FA). Briefly, applied an affine alignment of each diffusion-weighted image to b0 image to correct the eddy current distortions and motion artifacts. Accordingly, the b-matrix was reoriented based on the transformation matrix (Leemans and Jones, 2009). Then, the diffusion tensor elements were estimated (Basser et al., 1994) and the corresponding FA value of each voxel was calculated (Basser and Pierpaoli, 1996). All preprocessing procedures of the DTI data were performed with the FDT toolbox in FSL (http://www.fmrib.ox.ac.uk/fsl).

### Network construction

Nodes and edges are the two basic elements of a network. In this study, we constructed individual white matter (WM) structural networks using the following procedures.

#### Network node definition

The procedure of defining the nodes has been previously described (Bai et al., 2012; Cao et al., 2013; Zalesky et al., 2010) and was performed here using SPM8 software (http://www.fli.ion.ucl.ac.uk/spm). Briefly, individual T1-weighted images were coregistered to the b0 images in the DTI space. The transformed T1 were then nonlinearly transformed to the ICBM152 T1 template in the Montreal Neurological Institute (MNI) space. The inverse transformations were used to warp the automated anatomical labeling (AAL) atlas from the MNI space to the DTI native space. Discrete labeling values were preserved by using a nearest-neighbor interpolation method. By this procedure, we obtained 90 cortical and subcortical regions (45 for each hemisphere; **Table S1**), each representing a node of the network.

#### WM tractography

Diffusion tensor tractography was implemented with Diffusion Toolkit (http://www.trackvis.org/dtk/), using the “fiber assignment by continuous tracking (FACT)” method (Mori et al., 1999). All of the tracts in the dataset were computed by seeding each voxel with a FA that was greater than 0.2. Eight seeds were evenly distributed over the volume of the voxel. A streamline was started from each seed following the main diffusion direction from voxel to voxel, thus reconstructing WM fibers. The tractography was terminated if it turned an angle greater than 45 degrees or reached a voxel with a FA less than 0.2.

#### Network edge definition

For the network edges, two regions were considered structurally connected if there were at least one fiber streamline with two end-points that were located in these two regions (Bai et al., 2012; Shu et al., 2012; Zalesky et al., 2011). Specifically, we defined the number of interconnecting streamlines ended in two regions as the weights of the network edges. Therefore, for each participant, a weighted 90 × 90 WM structural network was constructed (**Fig S1**).

### Network analysis

All network analyses were performed with in-house software, Gretna (http://www.nitrc.org/projects/gretna/) (Wang, 2015 #425) and visualized by using BrainNet Viewer software (http://www.nitrc.org/progects/bnv/) (Xia, 2013 #426).

#### Network efficiency

To characterize the global topological organization of WM structural networks, we used the topological efficiency metrics: global efficiency (*E*_*glob*_) and local efficiency (*E*_*loc*_) (Rubinov and Sporns, 2010). The global efficiency of graph G measures the efficiency of the parallel information exchange in the network (Latora and Marchiori, 2001), which can be computed as:

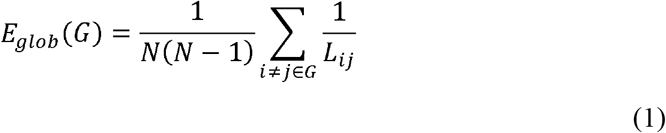

where *L*_*ij*_ is the shorted path length between node i and node j in G.

The local efficiency of G reveals how much the network is fault tolerant and shows how efficient the communication is among the immediate neighbors of the node i when it is removed. The local efficiency of a graph is defined as:

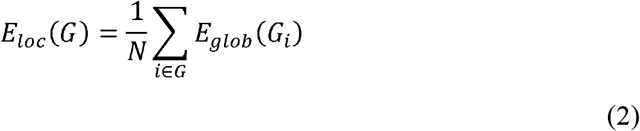

where *G*_*i*_ denotes the subgraph comprising all nodes that are immediate neighbors of the node i.

#### Regional nodal characteristics

To determine the nodal (regional) characteristics of the brain networks, we computed the nodal global and nodal local efficiency.

The nodal global efficiency, n*E*_*glob*_(*i*) is defined as (Achard and Bullmore, 2007):

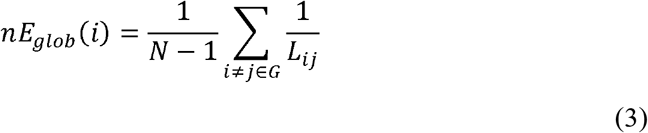

where *L*_*ij*_ is the shortest path length between node i and node j jn G. The nodal global efficiency measures the average shorted path length between a given node i and all the other nodes in the network.

The nodal local efficiency, *nE*_*loc*_(*i*) is defined as (Latora and Marchiori, 2001):

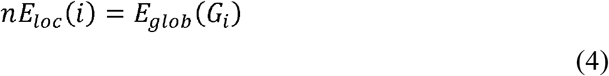

where *G*_*i*_ denotes the subgraph composed of the nearest neighbors of node i.

### Statistical analysis

To investigate the relationship between age and WM network metrics, partial correlation analyses were performed between age and WM network metrics within the participants, which removed the effects of sex and years of education. Then we carried out partial correlation analyses between WM network metrics and neuropsychological tests which removed the effects of age, sex and years of education. Finally we used the partial correlation analyses to investigate the relationship between WM network metrics and monthly mean change of neuropsychological tests. For regional properties, multiple comparisons were corrected by using the false discovery rate correction. All the above statistical analyses were performed by using statistical software (Matlab).

### Network-based predictive models

#### SVR training and evaluation

For predicting labels (monthly mean change of neuropsychological tests and baseline neuropsychological tests), we used support vector regression (SVR) with WM structure network metrics (nodal efficiency, nodal local efficiency) as features. Because of limited sample size of the dataset, leave-one-out cross-validation (LOOCV) was used to evaluate the SVR model. Each subject was designated as test subject in turns while the remaining ones were used to train the SVR predictor. The features of train sample and test sample were normalized with the mean and standard deviation of train samples. After that, partial correlation analyses were performed to select features correlating with labels. Then the residual of selected features were used to train the model. The procedure of SVR training and evaluation was shown in **Fig 1**. The hyper parameter C, epsilon and feature number were determined by grid searching. The performance of the model was quantified by the Pearson correlation coefficient between predicted values and labels. All the above analyses were performed based on the LIBSVM Toolbox (http://www.csie.ntu.edu.tw/~cjlin/libsvm/).

**Figure 1.**
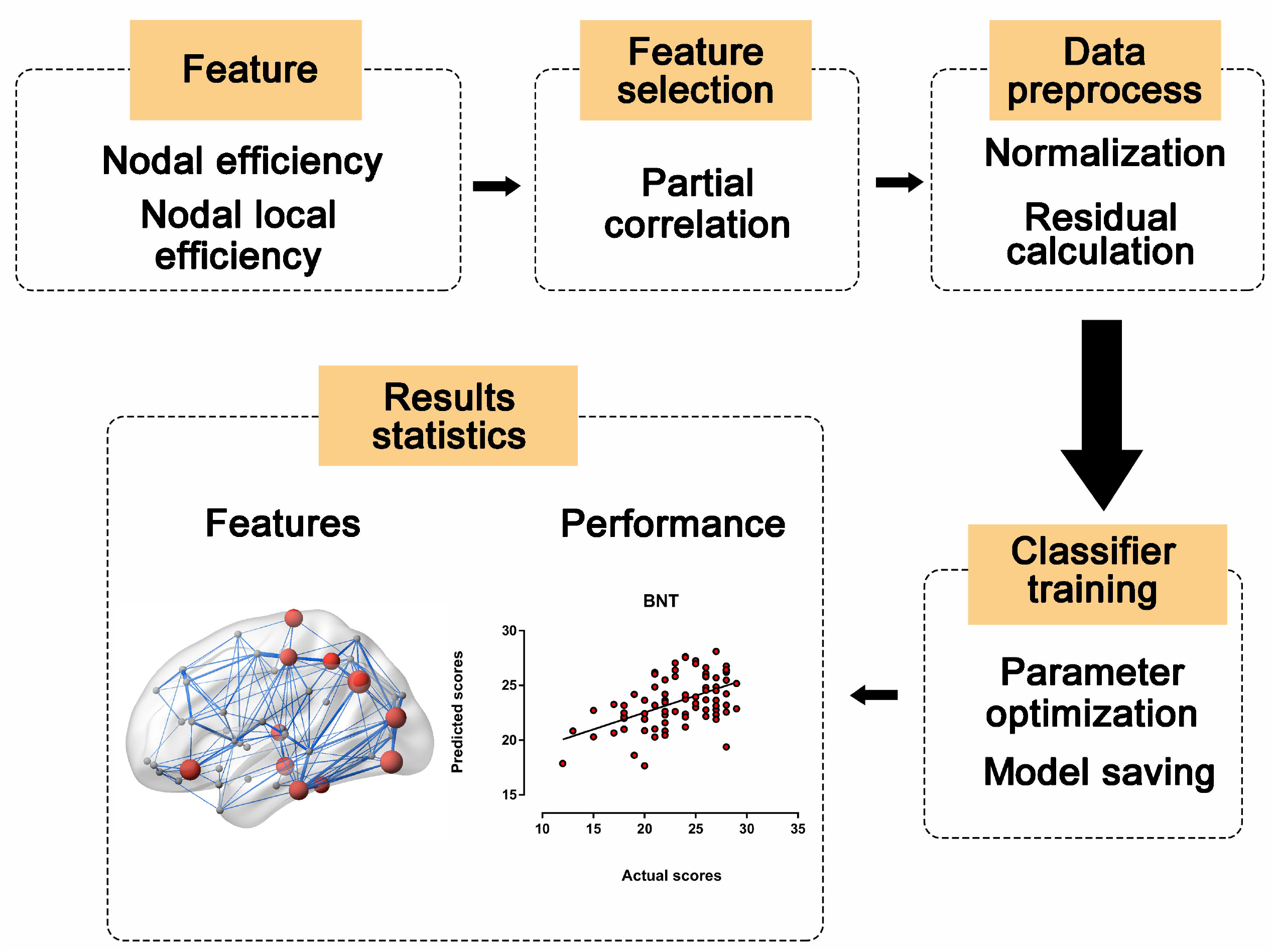
The procedure of prediction. 1) Used the nodal efficiency and nodal local efficiency as features for train the model. 2) Partial correlations were performed to select the features. 3) The features of test sample and train sample were normalized with the mean and standard deviation of train sample. After that, calculated the residual of features. 4) Trained the model and determined the optimized parameter by the grid searching. 5) Investigated the performance of model on the test sample and identified the most discriminative features.

#### Identification of the most important features

The essence of SVR model is determining a continuous function. The coefficients of the function quantify the importance to the prediction. Then the absolute coefficients of every fold of LOOCV were averaged as feature weights. The higher feature weights were determined to be, the more important of corresponding features. The weights of each brain region were obtained by summing the weights of the nodal efficiency and the nodal local efficiency of the corresponding brain region.

## Results

### Age effects on network metrics

Significant correlations were found between age and global network metrics (global efficiency: r=−0.29, p=0.007; local efficiency: r=−0.27, p=0.01). We identified regions with nodal efficiency correlated to age, which mostly located at right hemisphere (P < 0.05, false discovery rate corrected) (**Table 2, Fig 2**). The regions with nodal local efficiency correlated to age were not survived after false discovery rate correction.

**Table 1.**
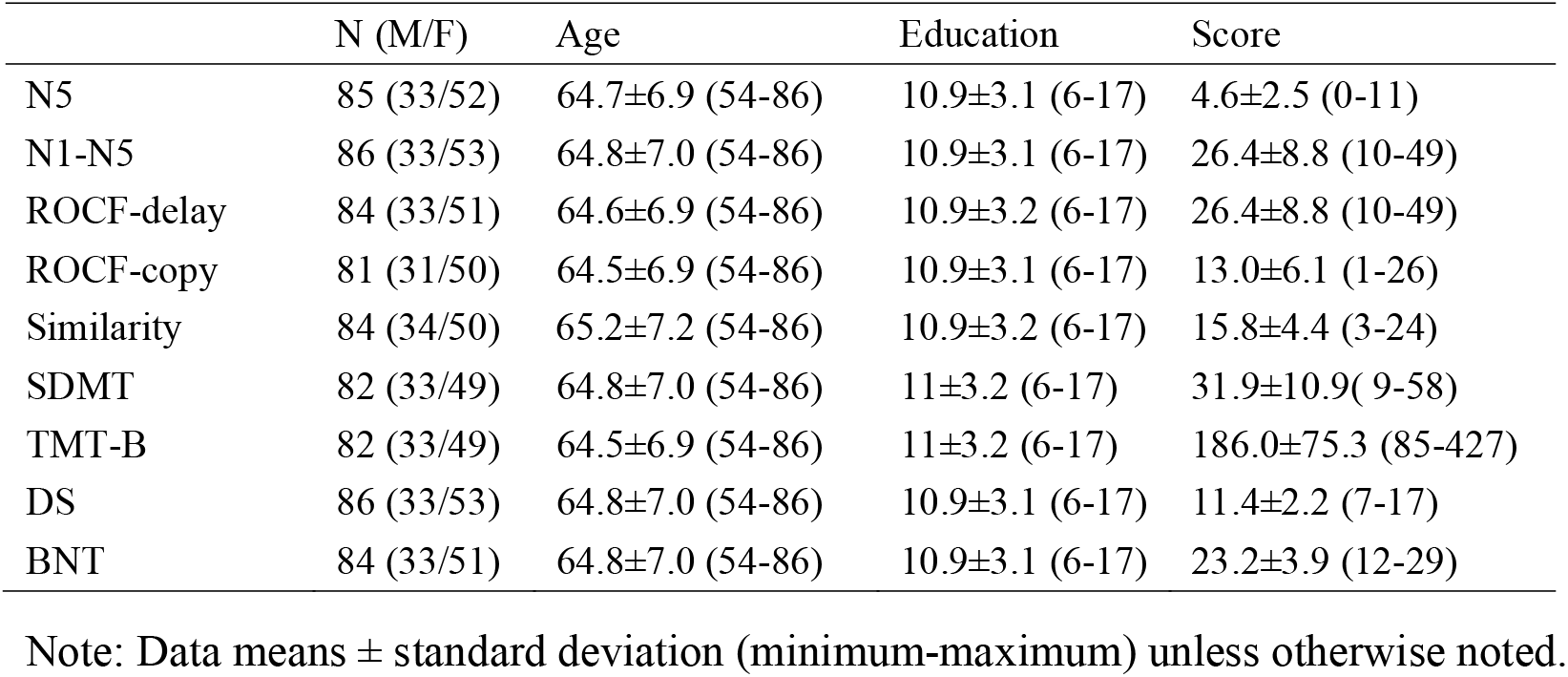
Neuropsychological characteristics of the sample.

|  | N (M/F) | Age | Education | Score |
| --- | --- | --- | --- | --- |
| N5 | 85 (33/52) | 64.7±6.9 (54-86) | 10.9±3.1 (6-17) | 4.6±2.5 (0-11) |
| N1-N5 | 86 (33/53) | 64.8±7.0 (54-86) | 10.9±3.1 (6-17) | 26.4±8.8 (10-49) |
| ROCF-delay | 84 (33/51) | 64.6±6.9 (54-86) | 10.9±3.2 (6-17) | 26.4±8.8 (10-49) |
| ROCF-copy | 81 (31/50) | 64.5±6.9 (54-86) | 10.9±3.1 (6-17) | 13.0±6.1 (1-26) |
| Similarity | 84 (34/50) | 65.2±7.2 (54-86) | 10.9±3.2 (6-17) | 15.8±4.4 (3-24) |
| SDMT | 82 (33/49) | 64.8±7.0 (54-86) | 11±3.2 (6-17) | 31.9±10.9( 9-58) |
| TMT-B | 82 (33/49) | 64.5±6.9 (54-86) | 11±3.2 (6-17) | 186.0±75.3 (85-427) |
| DS | 86 (33/53) | 64.8±7.0 (54-86) | 10.9±3.1 (6-17) | 11.4±2.2 (7-17) |
| BNT | 84 (33/51) | 64.8±7.0 (54-86) | 10.9±3.1 (6-17) | 23.2±3.9 (12-29) |

Note: Data means ± standard deviation (minimum-maximum) unless otherwise noted.

**Table 2.** Regions with nodal efficiency correlated to age.

| Regions | r | P | Corrected P |
| --- | --- | --- | --- |
| SMA.R | 0.41 | <0.001 | 0.007 |
| SMA.L | 0.41 | <0.001 | 0.004 |
| CAL.R | 0.40 | <0.001 | 0.004 |
| PCL.L | 0.39 | <0.001 | 0.003 |
| DCG.R | 0.38 | <0.001 | 0.005 |
| PCL.R | 0.38 | <0.001 | 0.004 |
| ORBsupmed.L | 0.37 | <0.001 | 0.005 |
| PCUN.R | 0.37 | <0.001 | 0.005 |
| PCUN.L | 0.35 | <0.001 | 0.007 |
| ORBsupmed.R | 0.35 | <0.001 | 0.008 |
| SFGmed.L | 0.33 | 0.002 | 0.013 |
| SPG.R | 0.32 | 0.002 | 0.016 |
| SFGmed.R | 0.32 | 0.002 | 0.015 |
| CAU.R | 0.32 | 0.003 | 0.017 |
| LING.R | 0.30 | 0.004 | 0.024 |
| SFGdor.R | 0.30 | 0.005 | 0.028 |
| ANG.R | 0.29 | 0.007 | 0.037 |
| PoCG.R | 0.28 | 0.007 | 0.036 |
| PreCG.R | 0.28 | 0.007 | 0.035 |
| CUN.R | 0.28 | 0.008 | 0.035 |

Note: L = left, R = right, SMA = Supplementary motor area, CAL= calcarine fissure and surrounding, PCL = paracentral lobule, DCG = median cingulate and paracingulate gyri, ORBsupmed = superior frontal gyrus, orbital part, PCUN = precuneus, SFGmed = Superior frontal gyrus, medial, SPG = Superior parietal gyrus, CAU = caudate nucleus, LING = lingual gyrus, SFGdor = Superior frontal gyrus, dorsolateral, ANG = Angular gyrus, PoCG = Postcentral gyrus, PreCG = Precentral gyres, CUN = cuneus.

**Figure 2.**
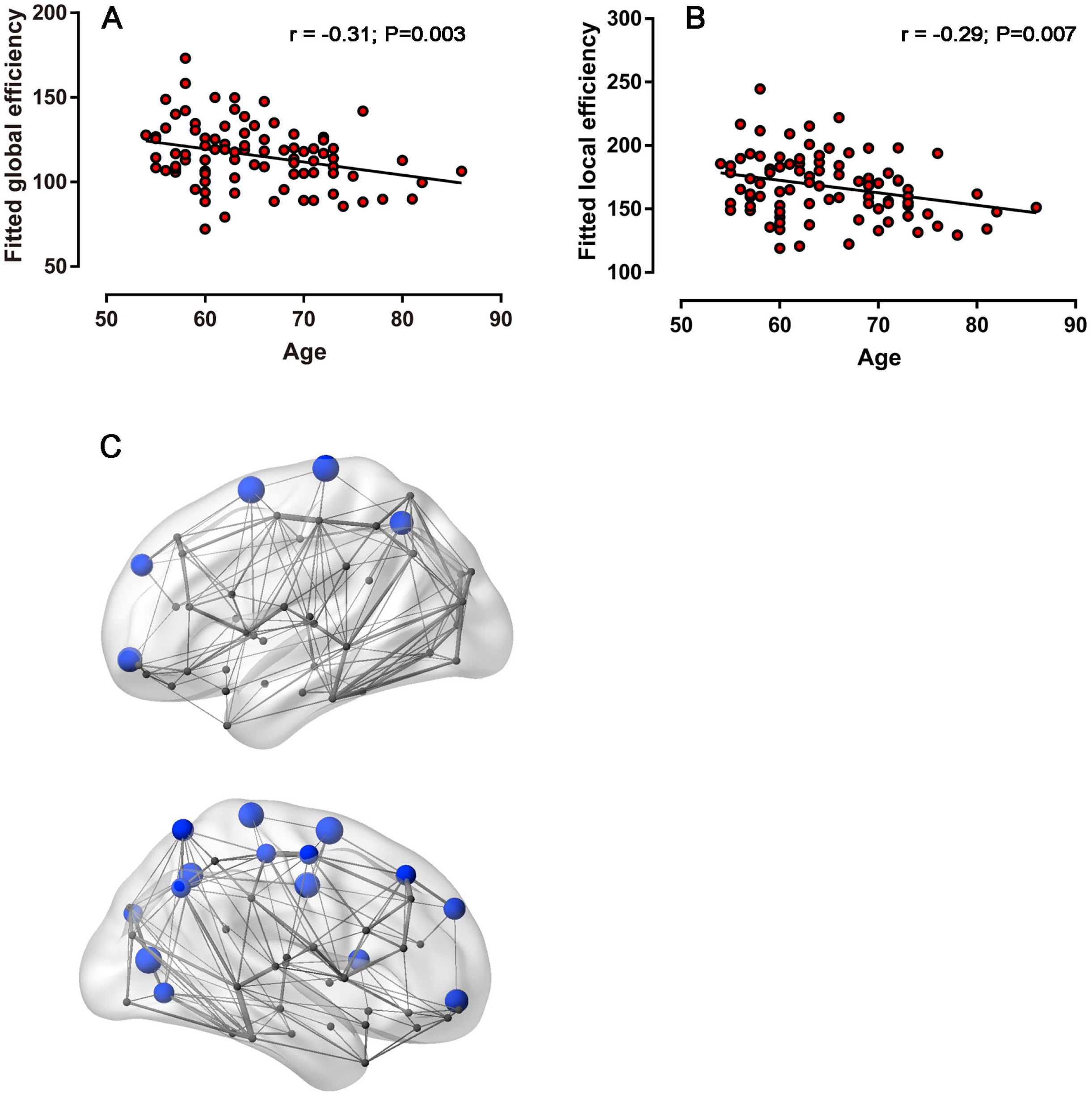
The correlation between age and network metrics. (A, B) Scatterplots show the significantly negative correlations between the age and the baseline WM brain network efficiency and local efficiency. (C) The regions with nodal efficiency correlated to age mostly locate…

### Relationship between network metrics and baseline cognitive function

Within the whole participants, network metrics correlated with baseline SDMT scores (global efficiency: r = 0.34, P = 0.002; local efficiency: r = 0.32, P = 0.003). The regions with nodal efficiency and nodal local efficiency correlated to SDMT were almost distributed the whole brain (P < 0.05, false discovery rate corrected) **(Fig 3**). There were no significant correlation between network metrics and other neuropsychological tests.

**Figure 3.**
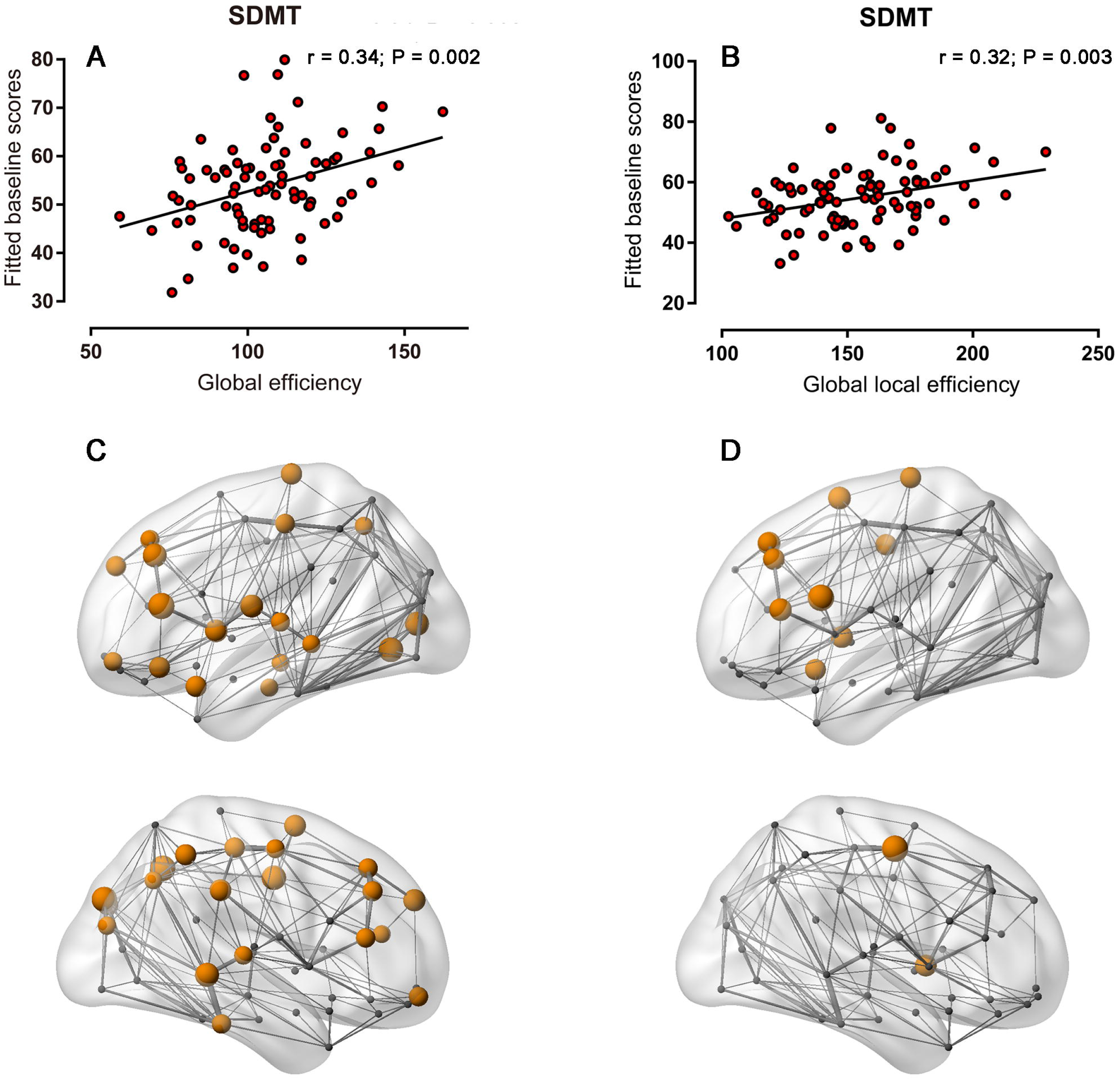
The correlation between baseline neuropsychological tests and network metrics. (A, B) The baseline SDMT scores are correlated to the baseline WM brain network efficiency and local efficiency significantly. The regions with baseline nodal efficiency correlated to baseline SDMT scores mostly locate in

### Prediction of baseline cognitive function

All the baseline scores of neuropsychological tests can be predicted apart from ROCF-delay recall. The predicted scores were correlated with the real scores (**Table 3**). The regions contributing to the prediction were specific for different neuropsychological tests (**Fig 4**). The top 5 regions of each model are listed in descending order of their weights (**Table 4**).

**Table 3.** The performance of SVR models

| Neuropsychological<br>test | Baseline |  | Monthly mean change |  |
| --- | --- | --- | --- | --- |
|  | r | P | r | P |
| N5 | 0.47 | < 0.001 | - | - |
| N1N5 | 0.46 | < 0.001 | - | - |
| ROCF-delay | - | - | - | - |
| ROCF-copy | 0.39 | < 0.001 | 0.38 | < 0.001 |
| Similarity | 0.38 | < 0.001 | 0.59 | < 0.001 |
| SDMT | 0.51 | < 0.001 | 0.57 | < 0.001 |
| TMT-B | 0.50 | < 0.001 | 0.43 | < 0.001 |
| DS | 0.35 | < 0.001 | 0.35 | < 0.001 |
| BNT | 0.52 | < 0.001 | 0.44 | < 0.001 |

**Table 4.** Top 5 regions showing the most important features for predicting baseline neuropsychological test.

| Neuropsychological test | Regions |
| --- | --- |
| N5 | DCG.R, PCUN.L, PAL.L, PCL.R |
| N1N5 | SOG.R, IOG.R, THA.R, PHG.L, PAL.L |
| ROCF-copy | PreCG.R, PAL.L, STG.R, IFGtriang.L, INS.R |
| Similarity | SOG.L, PCL.R, SMA.R, SMA.L, PAL.L |
| SDMT | SOG.R, CUN.L, MFG.R, PCL.L, STG.R |
| TMT-B | PUT.L, REC.R, IFGtriang.L, ORBmid.L, PoCG.R |
| DS | IFGtriang.R, IPL.R, PoCG.R, PCL.L, PAL.L |
| BNT | HES.R, SMA.L, MTG.R, PCL.L, ORBsupmed.R |

Note. Only 4 regions were used to predict the baseline N5 test.
L = left, R = right, DCG = median cingulate and paracingulate gyri, PCUN = precuneus, PAL = lenticular nucleus, pallidum, PCL = paracentral lobule, SOG = superior occipital gyrus, IOG = inferior occipital gyrus, THA = thalamus, PHG = parahippocampal gyrus, PreCG = precentral gyrus, STG = superior temporal gyrus, IFGtriang = Inferior frontal gyrus, triangular part, INS = insula, CUN = cuneus, MFG = middle frontal gyrus, PUT = lenticular nucleus, putamen, REC = gyrus rectus, ORBmid = middle frontal gyrus, orbital part, PoCG = postcentral gyrus, IPL = Inferior parietal, but supramarginal and angular gyri, HES = heschl gyrus, SMA = supplementary motor area, MTG = middle temporal gyrus, ORBsupmed = Superior frontal gyrus, medial orbital.

**Figure 4.**
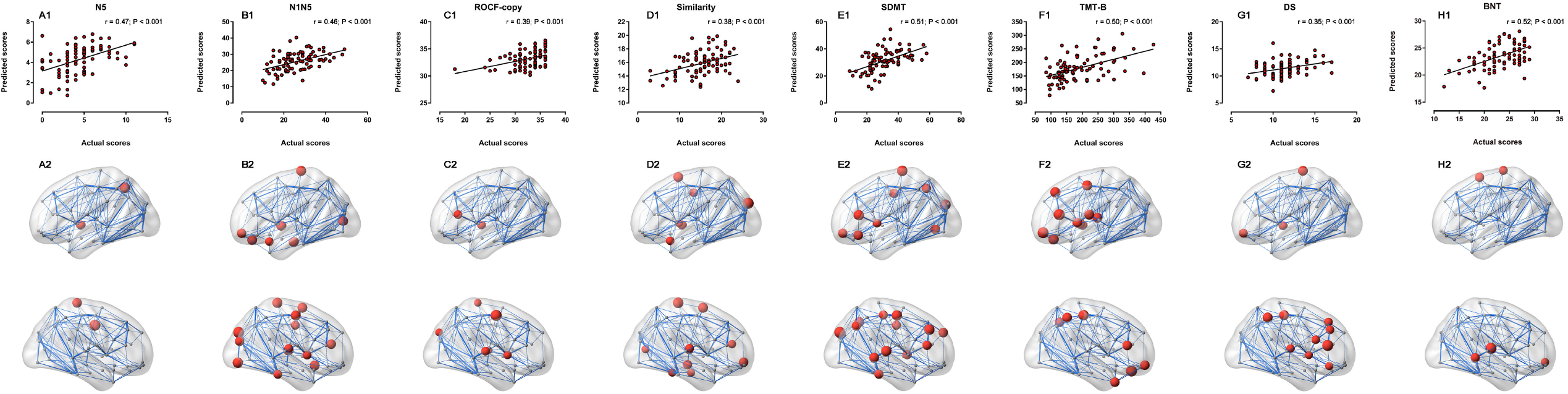
The correlation between the predicted baseline neuropsychological scores and the actual scores. (A1 - H1) The predicted values are correlated to the actual values significantly on the N5, N1N5, ROCF-copy, similarity, SDMT, TMT-B, DS and BNT tests. (A2 – H2) The red nodes are the regions contributing to the each baseline neuropsychological tests predictions correspondingly.

### Relationship between network metrics and cognitive change

Within the whole participants, network metrics correlated with monthly mean change of SDMT scores (global efficiency: r = 0.30, P = 0.007; local efficiency: r = 0.29, P = 0.01) and TMT-B scores (global efficiency: r =− 0.45, p < 0.001; local efficiency: r = −0.43, P < 0.001). The half regions with nodal efficiency correlated to monthly mean change of SDMT score located at occipital cortex (P < 0.05, false discovery rate corrected) (**Fig 5**) and no nodes with nodal local efficiency correlated to monthly mean change of SDMT score survived after false discovery rate correction. Almost the whole brain regions’ nodal efficiency correlated to monthly mean change of TMT-B score and the regions with nodal local efficiency correlated to monthly mean change of TMT-B score included left insula, left rolandic operculum, left superior temporal gyrus, left inferior temporal gyrus, left posterior cingulate gyrus, right precuneus and right putamen (P < 0.05, false discovery rate corrected) (**Fig 5**).

**Figure 5.**
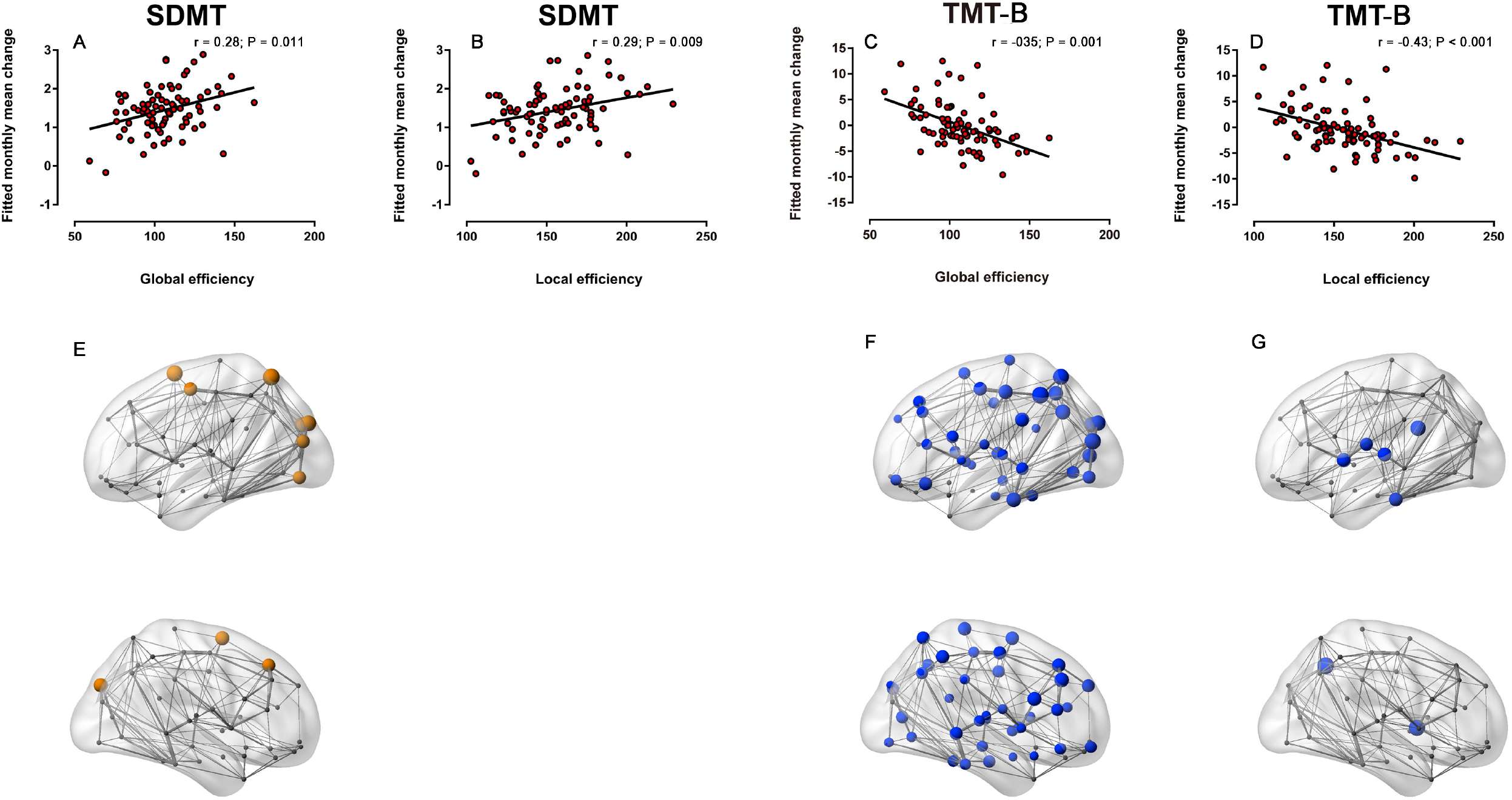
The correlation between monthly mean change of neuropsychological tests and network metrics. (A - D) The monthly mean change of SDMT and TMT-B scores are correlated to baseline network efficiency and local efficiency. (E) The regions with monthly mean change of SDMT scores correlated to baseline nodal efficiency mostly locate in occipital cortex. (F) Nearly all the brain regions’ baseline nodal efficiency are correlated to monthly mean change of TMT-B test. (G) The brain regions with baseline nodal local efficiency correlated to monthly mean change of TMT-B test are left insula, left rolandic operculum, left posterior cingulate gyrus, left superior temporal gyrus, left inferior temporal gyrus, right precuneus and right putamen.

### Prediction of cognitive change

All the monthly mean change of scores of neuropsychological tests can be predicted apart from ROCF-delay recall, N5 and N1N5. The predicted scores were correlated with the real scores (**Table 3**). The regions contributing to the prediction were specific for different neuropsychological tests and different from the prediction of baseline neuropsychological tests (**Fig 6**). The top 5 regions of each model are listed in descending order of their weights (**Table 5**).

**Table 5.** Top 5 regions showing the most important features for predicting monthly mean change of neuropsychological test.

| Neuropsychological test | Regions |
| --- | --- |
| ROCF-copy | OLF.L, MFG.R, SPG.R, THA.R, AMYG.L |
| Similarity | CAL.L, ORBsup.L |
| SDMT | MOG.L, SMA.L, SMA.R, SPG.L, SOG.L |
| TMT-B | MOG.L, PCUN.R, CUN.L, CAL.L, IPL.L |
| DS | SFGdor.L, REC.R, SMA.R, ORBinf.R, MOG.R |
| BNT | MFG.R, PreCG.R, MTG.R, ANG.L, IOG.L |

Note. Only 2 regions were used to predict the monthly mean change of similarity test.
L = left, R = right, OLF = olfactory cortex, MFG = middle frontal gyrus, SPG = superior parietal gyrus, THA = thalamus, AMYG = amygdala, CAL = calcarine fissure and surrounding, ORBsup = superior frontal gyrus, orbital part, MOG = middle occipital gyrus, SMA = supplementary motor area, SOG = superior occipital gyrus, PCUN = precuneus, CUN = cuneus, IPL = inferior parietal, but supramarginal and angular gyri, SFGdor = superior frontal gyrus, dorsolateral, REC = gyrus rectus, ORBinf = inferior frontal gyrus, orbital part, PreCG = precentral gyrus, MTG = middle temporal gyrus, ANG = angular gyrus, IOG = Inferior occipital gyrus.

**Figure 6.**
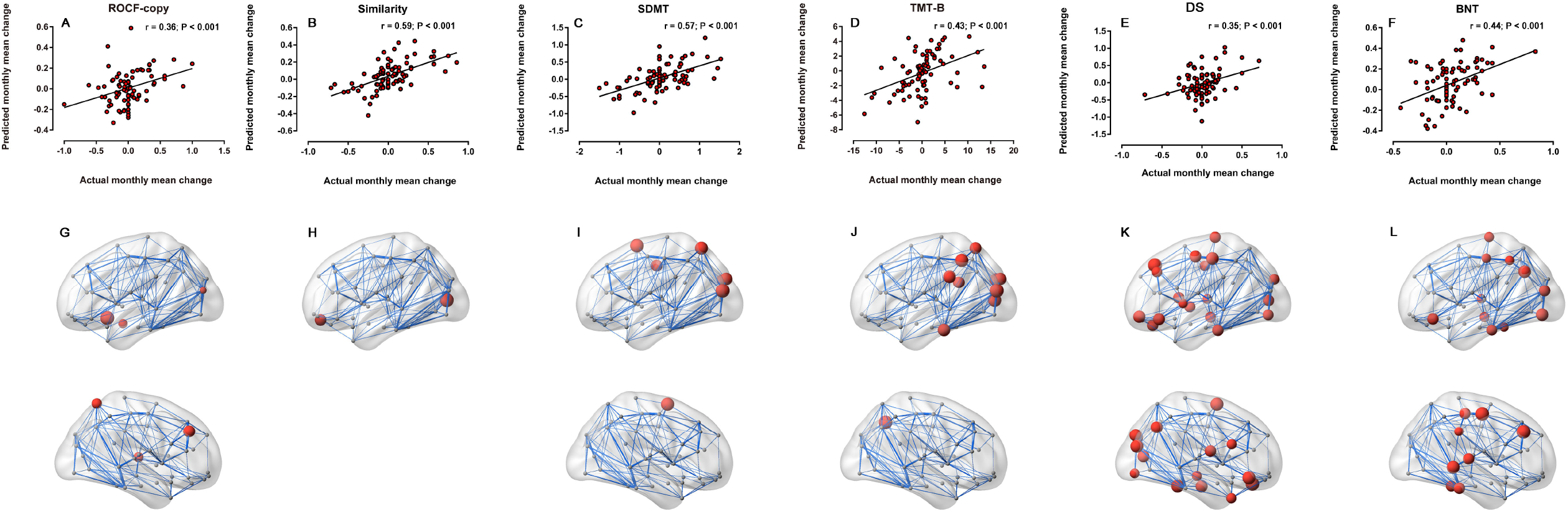
The performance of model of predicting monthly mean change of neuropsychological tests. (A - F) The predicted values are correlated to the actual values significantly on the ROCF-copy, similarity, SDMT, TMT-B, DS, and BNT tests. (G - L) The red nodes are the regions contributing to the each baseline neuropsychological test predictions correspondingly.

## Discussion

The present study demonstrated for the first time that WM network connectome parameters is able to predict longitudinal cognitive change over an 18-months observational period in older individuals. Baseline global and local efficiency were related with baseline performance of memory, executive functions, processing speed and reasoning, and only with significantly faster rate of decline in executive functions, and language independently of age, gender and education. More importantly, base on SUV analysis, predicted rate of cognitive decline was significantly related with actual rate of decline in multiple cognitive domains. These results suggested WM network connectome parameters are the strong predictor of longitudinal cognitive decline. Correlations between connectivity of specific regions and cognitive assessments at follow up were also observed. For example, the connectivity of right precuneus, left superior parietal gyrus, left posterior cingulate gyrus and so on were associated with the changed rate of executive function; the connectivity of left dorsolateral superior frontal gyrus, right gyrus rectus, supplementary motor area, inferior frontal gyrus and middle occipital gyrus.

The strong indicator of cognitive performance was the combined global and local efficiency of whole brain WM network, which was consistently related to a large spectrum of cognitive measures. The efficiency also predicted poor long-term cognitive decline during follow-up. The global efficiency represents the information transferring ability of WM tracts between remoter cortical regions. Generally, reduced global and local efficiency reflect a reduced efficiency of information transfer of WM networks. The most likely explanation for the observed associations between cognitive decline and the topological properties of WM network are the impaired capacity of information transfer across different regions of brain networks, which may be due to WM degeneration of both long- and short-range connections. The pathophysiological explanations of disconnection underlying diffusion changes include axonal degeneration and ischemic demyelination. The relationship with some possible factors of aging, such as cortical amyloid deposition, neural dysfunction, and vascular damage should be investigated in the future studies, in combination with some advanced multi-modal imaging techniques.

The finding that connectivity efficiency of WM network relates to cognitive decline suggests that the cognitive domains examined in the present study likely depend upon the coordination of multiple brain regions. Thus, the present findings are in line with “disconnection” theories of cognitive aging (Andrewshanna et al., 2007; Bartzokis et al., 2004; Bennett & Madden, 2014b; M et al., 2001) and suggest that reduced efficiency of WM network may disrupt communication across the whole brain, thereby contributing to cognitive decline. Several earlier cross-sectional studies have implied that diffusion changes are more sensitive to cognitive impairment than the conventional MRI findings(Vernooij et al., 2009; Xu et al., 2010). The topological efficiency was associated with cognitive functions, especially executive functions and processing speed (Lo et al., 2010) (Reijmer et al., 2013) (Shu et al., 2012). In the cognitively normal subjects, reduced WM microstructure is linked to poorer cognitive performance, and generally find stronger relationships between white matter microstructure and cognitive function(Madden et al., 2012).

Prospective longitudinal studies confirming the importance of disrupted white matter to cognitive decline have been rare. In a 1-year follow-up study of 35 patients with SVD, DTI correlated with executive functions more closely than other MRI parameters, but no relation was detected between their changes over time(Nitkunan, Barrick, Charlton, Clark, & Markus, 2008). Another study with 84 healthy middle-aged and elderly subjects has reported an association between DTI and working memory decline in 2-year follow-up, but executive functions and processing speed remained unchanged((Charlton, Schiavone, Barrick, Morris, & Markus, 2010)).Progression of reduced WM tracts predict cognitive decline, with the most pronounced effects on processing speed and executive functions (Jokinen et al., 2011; Kramer et al., 2007; Prins et al., 2005; van Dijk et al., 2008). Meanwhile, other studies found the WM tracts predict the episodic memory over time, but not executive function (Fjell et al., 2016; Rabin et al., 2018). Abnormal diffusion changes in the WM tracts also show the strong predictive value for the discrimination between MCI and cognitively normal subjects(Chua et al., 2009; Rose et al., 2006; Sexton, Kalu, Filippini, Mackay, & Ebmeier, 2011) and the conversion from MCI to AD within 1.5 years(Fellgiebel et al., 2006). Previous studies suggest the WM show the strong predictive value for cognitive decline. Our study was the first study about the predictive value of efficiency of WM network.

We examined for the first time not only the correlations of neuroanatomical connections as a system, but also whether neuroanatomical connectivity can be segregated to explain specific cognitive decline. Because particular brain regions are important for specific functions, the capacity of information flow within and between regions is also crucial. Our results found that the efficiency of WM network had an influence on the changes of executive function, working memory, attention, language and memory. The predictors of cognitive function are important because impairment in this domain increases the likelihood of conversion to dementia (Cloutier, Chertkow, Kergoat, Gauthier, & Belleville, 2014; Ewers et al., 2014).Correlations between connectivity of specific regions and cognitive assessments were also observed.

The regional distribution of local networks associated with the executive domain included right precuneus, left superior parietal gyrus, left posterior cingulate gyrus and so on. The connectivity efficiency of these regions has the stronger predictive value for executive function decline than other cognitive functions. Previous studies have observed a more apparent relationship between white matter integrity and executive function compared to that with memory(Chang et al., 2015; Hedden et al., 2012; Nir et al., 2013). Some DTI network analyses find that decreased nodal or global efficiency correlates with executive function or working memory (Fischer, Wolf, Scheurich, & Fellgiebel, 2015; Kim et al., 2016). These tasks are largely dependent on fluid intelligence and may not reflect a unique aspect of functioning, which is thought to be more vulnerable to damage to cortical connectivity, in contrast to crystallized intelligence(Ritchie et al., 2015; Roca et al., 2010). In agreement with previous studies of older adults with cerebral small vessel disease (Jacobs et al., 2013, Association between white matter microstructure, executive functions, and processing speed in older adults: the impact of vascular health. Hum Brain Mapp. 34:77–95.; Ciulli et al., 2016), the results of our comprehensive the whole brain network analyses, suggest that poorer white matter microstructure throughout the cortex, not exclusively frontal associated tracts, may serve as an effective indicator of executive function performance. Similar to the executive function, working memory, which also depended on the multiple cognitive domains, has been consistently associated with the efficiency of frontal, parietal and occipital regions. It is therefore not surprising that poorer connectivity both globally in the brain and locally in large numbers of regional cortical areas could result in poorer performance. Working memory is very important to cognitive aging. In elderly persons, the grey matter atrophy, decreased functional connectivity and reduced white matter integrity especially in frontal and parietal regions appeared. The decreased efficiency of WM network with age is sensitive to the deficits of working memory.

The strongest predicted value for cognitive decline of attention located in parietal and occipital regions, such as superior parietal gyrus, middle occipital gyrus, supplementary motor area and so on. The superior parietal gyrus is related with visuospatial and attentional processing(Nachev & Husain, 2006). It receives a great deal of visual and sensory input, and also associated with deficits on the manipulation and rearrangement of information(Koenigs, Barbey, Postle, & Grafman, 2009). The test used to measure attention in this study, Symbol digital modalities test, places demands on attention shift and perceptual reasoning, in addition to more basic visuospatial perception (??). Thus our finding of multiple local efficiency networks represented several parietal and occipital regions which were consistent with the multifactorial nature of this test. The parietal and frontal regions are also the effective predicted index for the changed rate of spatial-visual ability. The fronto-parietal regions have been identified for visuospatial transformations(Goebel, Linden, Lanfermann, Zanella, & Singer, 1998) (Carpenter, Just, Keller, Eddy, & Thulborn, 1999), visual search (Leonards, Sunaert, Hecke, & Orban, 2000), and spatial memory (Diwadkar, Carpenter, & Just, 2000) . We could have expected to see significant findings in fronto-parietal regions for the figure copy test.

In our study, we found that the temporal and parietal regions that were significant were consistent with the findings from studies looking at functional and structural correlates of these language tasks (Apostolova et al., 2008; Grossman et al., 2004; Whitney et al., 2014) . We could have expected to see significant findings in the middle frontal region and middle and inferior temporal regions (Pihlajamaki et al., 2000). However, memory may not be related to the same extent as executive to corticocortical connectivity because of the key role some subcortical structures such as the entorhinal cortex and hippocampus play in this cognitive domain (Raz et al., 2005). Entorhinal cortex is considered as the key region for age related cognitive decline in healthy aging. However, the connectivity between the entorhinal cortex and other subcortical structures with cortical regions may not appear in our study.

Strengths of this study include 1) the longitudinal design, with annual testing for up to 2 years; 2) the assessment of multiple cognitive domains; 3) the relatively large sample size (n = 100), and 4) the whole brain white matter networks examined.

## Conclusions

We present novel findings on the prognostic value of WM network in predicting long-term cognitive outcome in a large cohort of elderly healthy subjects. In particular, the loss of WM connectivity as detected by DTI heralds a faster rate of decline of executive functions and processing speed. Moreover, DTI network changes in elderly subjects predict poor prognosis in terms of cognitive disability. These results provide new insight into the clinical validity and sensitivity of DTI as an early marker for cognitive decline. Our study highlight the need for a better understanding of the ways in which white matter microstructural alterations can be prevented or ameliorated in normal aging. Importantly, the benefits of the WM network are the availability in clinical stings, very short acquisition times, and less demands for equipment. It is also a well-established technique and provides more information that a regional assessment.

