## Supplementary figures and images for "The white matter network in cognitive normal elderly predict the rate of cognitive decline"

### Supplemental Figure 1

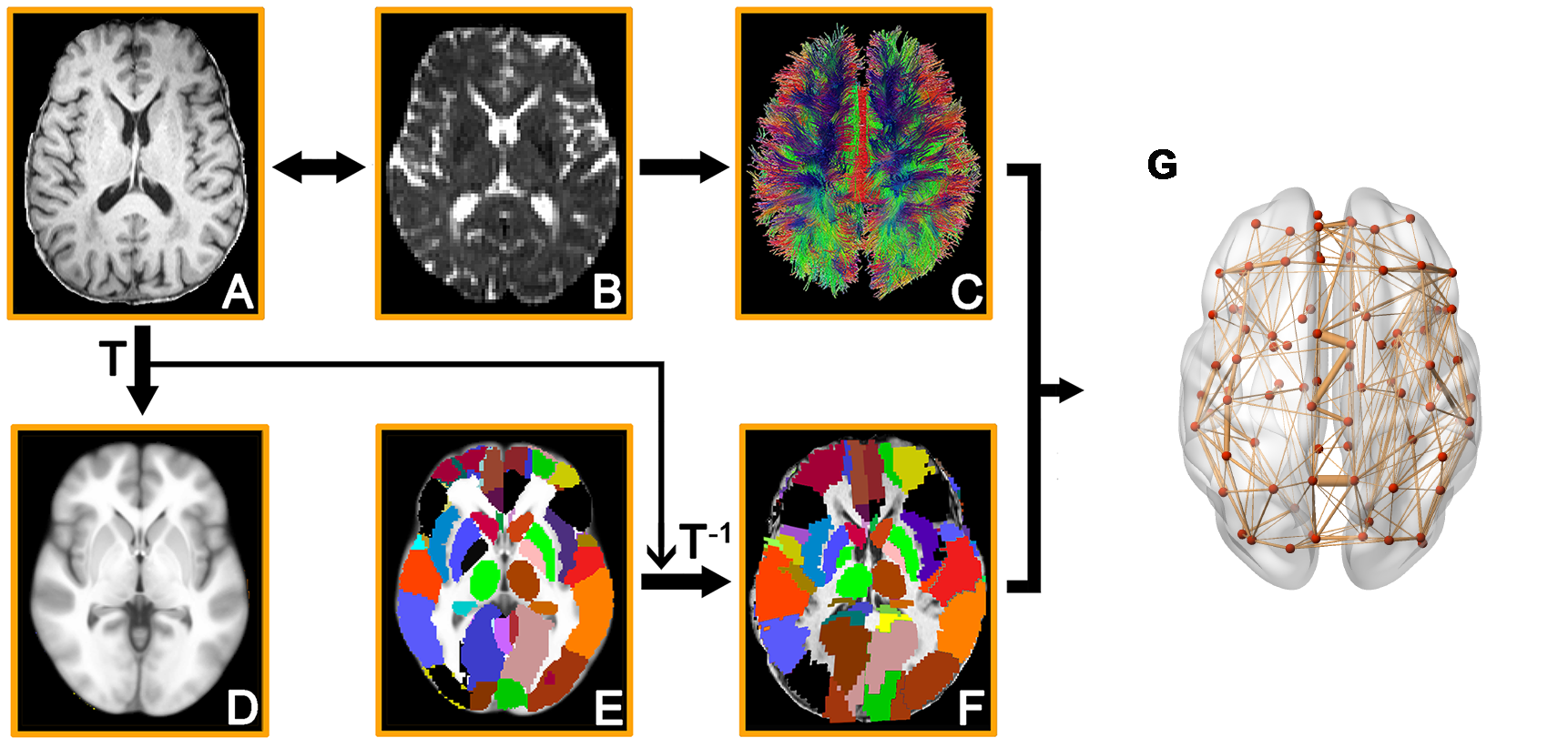
